# Molecular Ecology of Coral Reef Microorganisms in the Western Indian Ocean coast of Kenya

**DOI:** 10.1101/2020.03.04.976431

**Authors:** Sammy Wambua, Hadrien Gourlé, Etienne de Villiers, Oskar Karlsson, Nina Wambiji, Angus Macdonald, Erik Bongcam-Rudloff, Santie de Villiers

## Abstract

Coral reefs face increased environmental threats from anthropomorphic climate change and pollution, from agriculture, industries and tourism. They are economically vital for many people worldwide, and harbour a fantastically diverse ecosystem, being the home for many species of fish and algae. Surprisingly little is known about the microbial communities living in and in the surrounding of coral reefs. Here we employ high throughput sequencing for investigating the bacteria living in the water column and upper sediment layer in close proximity to coral reefs on the Kenyan coast of the West Indian Ocean. We show that while the read-level taxonomic distribution of bacteria is similar with ones obtained from 16S metabarcoding, whole metagenome sequencing provides valuable functional insights not available with 16S metabarcoding. We find evidence of pollution, marked by the presence of *Vibrio* and more importantly the presence of antibiotic resistance notably to vancomycin, that we attribute to the use of avoparcin in agriculture. Additionally, 175 bacterial genomes not previously sequenced were discovered.

Our study is the first whole-metagenome study from the West Indian Ocean, provides a much-needed baseline to study microbes surrounding coral reefs under different conditions as well as the microbiome of coral reefs.

## Introduction

Coral reefs are one of the most biodiverse ecosystems in the world, thereby providing vast ecological and socio-economic resources. However, coral reefs and consequently their invaluable ecosystem services are increasingly under threat globally due to climate change and a range of other human-related pressures, such as pollution from agriculture, industries and tourism-related activities. These stressors lead to coral degradation the onset of which is most often marked by bleaching following the expulsion of the symbiotic algae.

Addressing the challenges leading to coral death requires a comprehensive understanding of corals and their interactions with other members of the reef ecosystems. Appreciation of this fact has led to increased interest in marine microbiology research as microorganisms are thought to be critical for reef ecosystem processes including coral homeostasis, nutrition and protection against disease (Godoy-Vitorino, Ruiz-Diaz, Rivera-Seda, Ramírez-Lugo, & Toledo-Hernández, 2017). Microbial communities are also known to respond and adapt quickly to disturbance (Ainsworth, Thurber, & Gates, 2010). Therefore, studying coral reef-associated microorganisms, as well as microorganisms living in close proximity of coral reefs, holds the potential to help in improving the capacity to predict responses of coral reef ecosystems to changing environmental conditions.

The advancement of DNA sequencing and analysis technologies, allowing sequencing DNA directly without the need of cultivating organisms in laboratory settings, has augmented our understanding of the complexity and diversity of natural microbial populations (Biller et al., 2018). Most oceanic and costal surveys to date have used 16S metabarcoding, which provides precious insights about population complexity and diversity, but only a broad overview of the taxonomic distribution across samples with (i) limited resolution, especially for poorly characterised samples such as environmental ones and (ii), no or poor functional and metabolic insights of the sequenced communities (Poretsky, Rodriguez-R, Luo, Tsementzi, & Konstantinidis, 2014; Rausch et al., 2019). Whole Metagenome sequencing (WMS) on the other hand, has the potential for better taxonomic resolution, theoretically at the species level, even though as for 16S taxonomic classification is highly tied to the database used and the type and provenance of samples. Additionally, WMS also allows for reconstructing draft genomes from metagenomes, providing exceptional phylogenomic insights as well as opening up potential for functional annotation. Combined with recent advances in protein assembly directly from metagenomic data, the method shows great promise to harness metabolic pathways and functional knowledge from microbial communities.

The Western Indian Ocean (WIO), however, remains the least studied (Díez et al., 2016) of the oceans, despite hosting the second largest hotspot for coral biodiversity globally, partly due to the region’s lack of technological capacity. Marine metagenomes, like most environmental metagenomes, are diverse, making them especially difficult and expensive to analyse and interpret. Here we deeply sequence microbiomes from the water column and the upper sediment layer of three coastal reefs in Kenya, in an attempt to give an excellent taxonomic overview of WIO coastal communities and to unravel the functional characteristics of those rich environments.

## Methods

### Study Sites

The study was conducted at three marine protected areas (MPA) covering three of the five counties on the coast of Kenya Indian Ocean (Fig 1). Consisting of fringing reefs, each of the three sites was selected for its distinct human activities. Located about 118 km north of Mombasa, Malindi Marine National Park and Reserve is the oldest MPA in Kenya, having been gazetted in 1968 (McClanahan, Kaunda-Arara, & Omukoto, 2010). Sampling was done close to the reserve (3°15’35.1”S, 40°08’40.0”E) where artisanal fishing is allowed. The marine park is famous for glass-bottom boat tours and snorkelling among other recreational touristic activities. It also experiences significant year-round discharge of freshwater and sediments from the Sabaki River which runs through a catchment area dominated with agricultural settlements (Munyao, Tole, & Jungerius, 2003; van Katwijk et al., 1993). Mombasa Marine National Park and Reserve (3°59’45.7”S, 39°44’50.1”E) was established in 1986 (Ngugi, 2001)– with restrictions of protection being enforced commencing 1991 (Tuda & Omar, 2012) – and is, arguably, the most visited of Kenya’s marine parks by both local and international tourists (Owens, 1978). Due to its proximity to the urbanised touristic city, the park experiences pollution from hotels, hospitals, domestic and industrial waste disposals (Okuku et al., 2011; Stephen Mwangi, David Kirugara, Melckzedeck Osore, Joyce Njoya, Abdalla Yobe and Thomas Dzeha, 2001). Moreover, because Mombasa is the primary port serving inland eastern and central African countries, the park is also impacted by marine traffic activities including oil pollution and dredge-spoil dumping (Stephen Mwangi, David Kirugara, Melckzedeck Osore, Joyce Njoya, Abdalla Yobe and Thomas Dzeha, 2001). Lastly on the southern coast, 90 km from Mombasa, the Kisite-Mpunguti site (4°42’54.0”S, 39°22’23.8”E) is a protected area that comprises of Kisite Marine National Park and Mpunguti Reserve created in 1973 and gazetted in 1978. It is bordered by sparsely populated coral islands and experiences the least human activities because it is 11 km offshore (Emerton & Tessema, 2001).

**Figure1:**
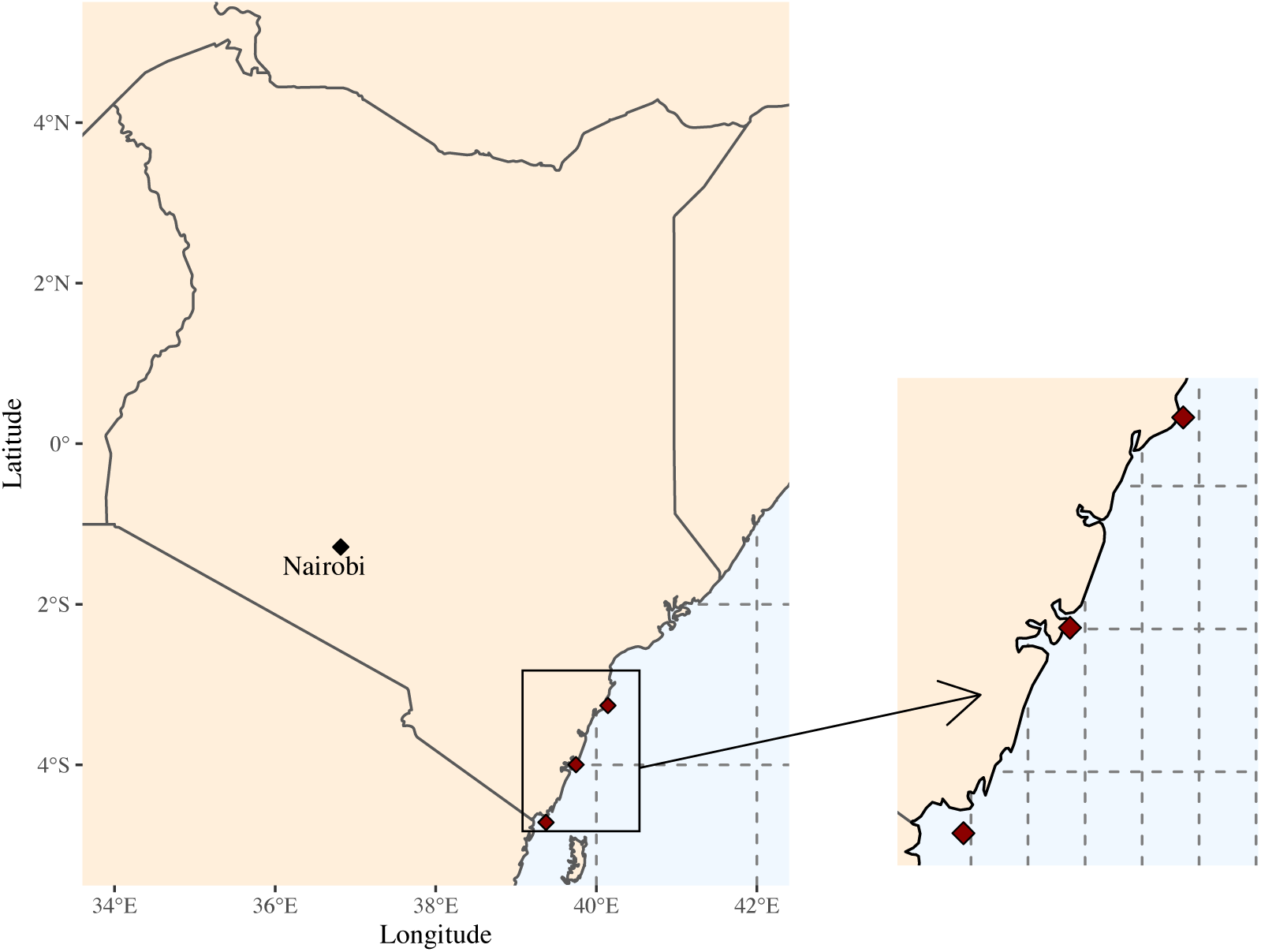
Location of the sampling sites. The three sampling sites are indicated in red, respectively from top to bottom: Malindi, Mombasa and Kisite. Each of the three sites was selected for its human activities. Sampling was done in 2016 and 2017, 200 - 500 m from the shore, at a depth of 1-2 m during low tides in the morning hours.

### Field Procedures

Sampling was done between 2016 and 2017 within the coral reefs, 200 - 500 m from the shore, at a depth of 1-2 m during low tides in the morning hours. At each site, seawater was collected in 5 L water bottles within 10 - 20 cm of a colony of *Acropora* spp. - the dominant coral species at the sites - for microbial DNA isolation. Additional triplicate 50 mL seawater samples were collected in disposable centrifuge tubes for nutrient analysis. A 10 mL syringe barrel was used to collect 2 cm column of sediment at the base of each sampled coral colony, 0.25 g of which was suspended in a bead tube containing inhibitor-dissolving and nucleic acid-preserving buffer. Samples were transported on ice to the laboratory for processing, typically within 3 hours of sampling.

Physicochemical parameters including water temperature, pH, salinity, and dissolved oxygen were determined *in situ*, at the time of sampling within the coral reefs, using portable multiprobe water quality meters, per manufacturer’s instructions (YSI Inc., Yellow Spring, OH).

### Laboratory Procedures

#### Nutrient testing

Spectrophotometry methods were used to determine seawater concentrations of nitrate (NO_2_^-^ -N), nitrite (NO_3_^-^ -N), ammonium (NH_4_^+^ -N) and phosphate (PO_4_^3-^ -P) nutrients (Supplementary Table 1 and 2) as described in Ongore et al., 2013 (Ongore et al., 2013).

#### DNA isolation

For each site, 4 L of seawater was vacuum-filtered (VWR, West Chester, PA, USA) through a 0.2-um pore size membrane (Pall Corporation, Port Washington, NY, USA) to capture microbial cells which were then added to a bead tube with lysis buffer. PowerWater ® DNA isolation kits were used to isolate microbial DNA from seawater samples while sediment samples were extracted with PowerSoil^®^ DNA isolation kits according to the manufacturer’s instructions (Mo Bio, Inc., Carlsbad, CA, USA). Quality and quantity of DNA were checked by Nanodrop and suitability for sequencing of DNA samples for metagenomics analysis was confirmed with 1% agarose gel (Rohwer, Seguritan, Azam, & Knowlton, 2002).

#### Library preparation and DNA sequencing

Sequencing libraries were prepared from 1µg of DNA according to the manufacturers’ preparation guide # 15036187 using the TruSeq DNA PCR-free library preparation kit (20015962/3, Illumina Inc.).

Briefly, the DNA was fragmented using a Covaris E220 system, aiming at 350bp fragments. Resulting DNA fragments were end-repaired, and the 3’ end adenylated to generate an overhang. Adapter sequences were ligated to the fragments via the A-overhang and the generated sequencing library was purified using AMPure XP beads (Beckman Coulter). The quality of the library was evaluated using the FragmentAnalyzer system and a DNF-910 kit. The adapter-ligated fragments were quantified by qPCR using the Library quantification kit for Illumina (KAPA Biosystems/Roche) on a CFX384Touch instrument (BioRad) before cluster generation and sequencing.

A 200 pM solution of the individual sequencing libraries was subjected to cluster generation and paired-end sequencing with 150bp read length using an S2 flowcell on the NovaSeq system (Illumina Inc.) using the v1 chemistry according to the manufacturer’s protocols.

Base-calling was done on the instrument by RTA 3.3.3 and the resulting .bcl files were demultiplexed and converted to fastq format with tools provided by Illumina Inc., allowing for one mismatch in the index sequence.

Sequencing was performed by the NGI SNP&SEQ Technology Platform in Uppsala, Sweden www.sequencing.se.

### Bioinformatics Analyses

The raw Illumina reads were trimmed at Q5 threshold (Macmanes, 2014), and the adapters were removed using fastp v0.19.5 (Chen, Zhou, Chen, & Gu, 2018). Trimmed sequences were deposited to the European Nucleotide Archive under the study accession PRJEB30838.

The trimmed reads were assigned a taxonomic classification using a combination of Kraken v2.0.8 and bracken v2.2 against the nt database using default parameters (Lu, Breitwieser, Thielen, & Salzberg, 2017; Wood, Lu, & Langmead, 2019). Rarefaction curves were computed using R and vegan v2.5 (Oksanen et al., 2019; R Core Team, 2018).

The samples were assembled using megahit v1.1.4 with the options --k-min 27 --k-max 147 -- k-step 10 (Li, Liu, Luo, Sadakane, & Lam, 2015). The reads were then mapped to the assemblies with bowtie v2.2.9 (Langmead & Salzberg, 2012) using default parameters and binned into draft genomes with metabat v2.11.1 with option –minContig 1500 (Kang et al., 2019). The genomes bins were then quality checked and refined with checkm v1.0.7 and refinem v0.0.24 (Parks, Imelfort, Skennerton, Hugenholtz, & Tyson, 2015; Parks et al., 2017) (scripts and refining parameters are available at https://osf.io/5fzqu/), and the best genome bins were annotated using prokka v1.10 (Seemann, 2014) and eggnog-mapper v1.0.3 (Huerta-Cepas et al., 2017), as well as placed phylogenetically using gtdbtk v0.3.2 (Chaumeil, Mussig, Hugenholtz, & Parks, 2019). The tree figure was generated using ggtree v1.14.6 (Yu, Smith, Zhu, Guan, & Lam, 2017).

The trimmed reads were also assembled directly at the protein level using plass (commit 26b5d6625a2fbef4cfaab4bfaa99b1682d35921c) (Steinegger, Mirdita, & Söding, 2018). The resulting assemblies were then clustered at 40, 50 and 90% identity using cd-hit v4.7 (Fu, Niu, Zhu, Wu, & Li, 2012) and annotated using eggnog-mapper v1.0.3.

## Results

### Taxonomic distribution of sediment and water associated organisms

Of the 4.2 billion reads obtained, 608 million were classified as bacteria, 16 million as archaea, 12 million as viruses, 537 million as eukaryotic sequences and 3 billion remained unclassified. The bacterial species richness was estimated at around 15000 for all samples (Supplementary Fig 1), which is high but within reasonable magnitudes according to previously published large metagenomes (Rodriguez-R & Konstantinidis, 2014). The vast majority of bacterial sequences were identified as proteobacteria (Figure 2). The second most abundant phyla were *Bacteroidetes* in the water samples and *Cyanobacteria* in the sediment samples.

**Figure 2:**
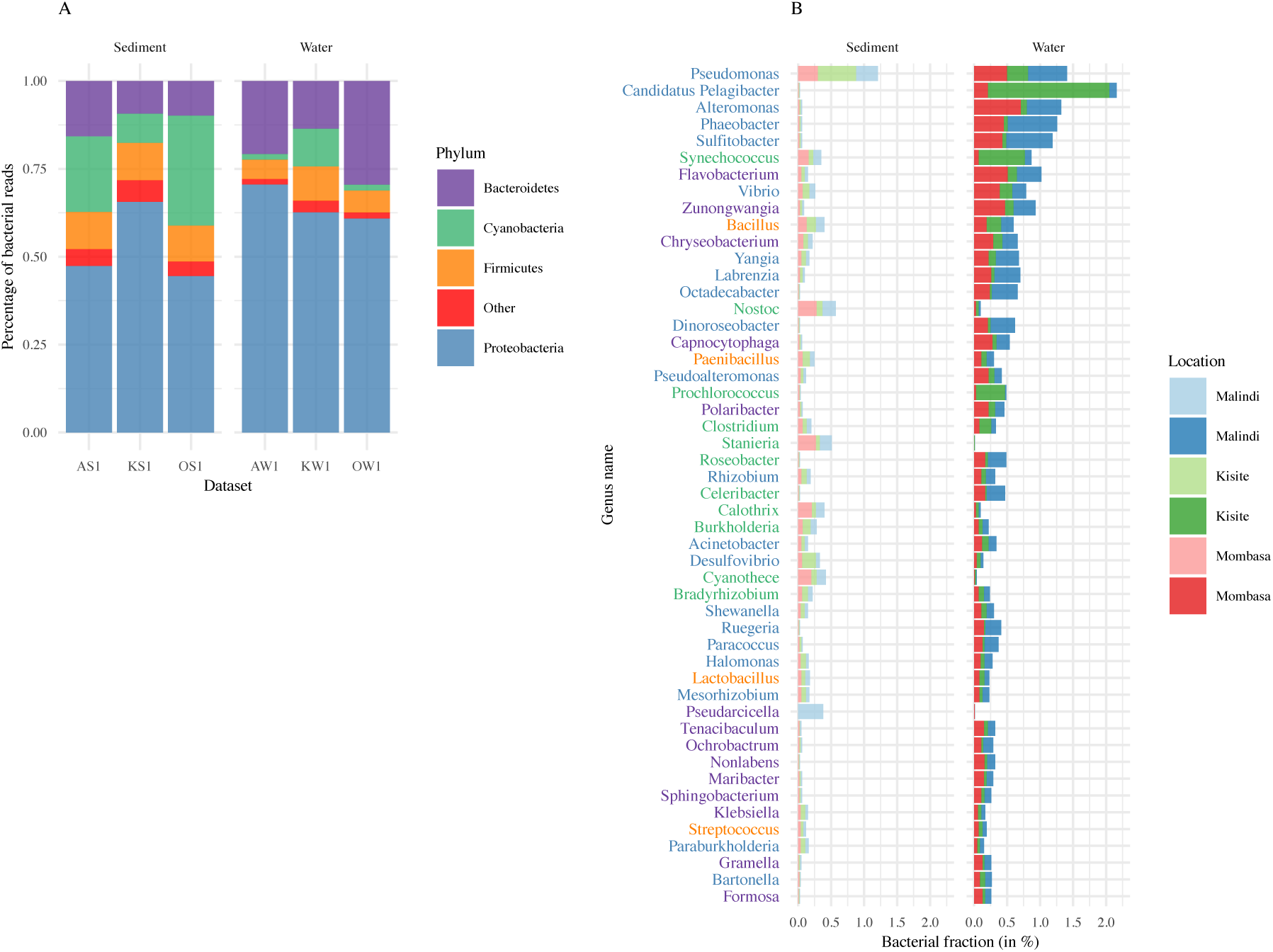
Taxonomic composition of the samples. Panel A shows the four most abundant bacterial phyla in each samples. Panel B shows the distribution of the 50 most abundant genera in the sediment and water samples. The majority of classified bacterial sequences were Proteobacteria.

At the genera level, *Pseudomonas* was ubiquitous in all samples (Figure 2), and dominant in the sediment, whilst the diversity of Proteobacteria was found to be much higher in the water samples, constituting the five most abundant genera. A non-negligible fraction of *Vibrio* was also found in all sampling sites, mainly in the water samples. The most abundant cyanobacteria in the Sediment samples were *Nostoc, Stanieria, Cyanothece, Calothrix* and *Synechococcus* (with *Synechococcus* being also found in abundance in the water samples). In water, Bacteroidetes were dominated by Flavobacteriaceae, more specifically the genera *Flavobacterium, Zunongwangia, Chryseobacterium* and *Capnocytophaga*. Lastly, abundant traces of *Bacillus, Paenibacillus, Lactobacillus* and *Streptococcus* were found.

The classified archaeal reads were mostly divided into two phyla: Euryarchaeota and Thaumarchaeota, with the former dominating the water samples and the latter the sediment samples. The Thaumarchaeota fraction is explained by the presence of *Nitrosopumilus*, a common organism living in seawater. The Euryarchaeota fraction is a bit more diverse but is mainly comprised of methanogens (Supplementary Fig 2).

#### Assembly and Functional annotation of proteins

A total of 21 million proteins were assembled, 12 million of which were unique proteins (after clustering at 90% identity). A total of 424 distinct KEGG pathways were identified in the data, which supports high diversity in function, especially in the water samples. Of note is that, in the 20 most abundant pathways, we found a total of 952017 genes associated with antibiotic biosynthesis, making it the second most abundant KEGG category after Biosynthesis of secondary metabolites (Fig. 3).

**Figure 3:**
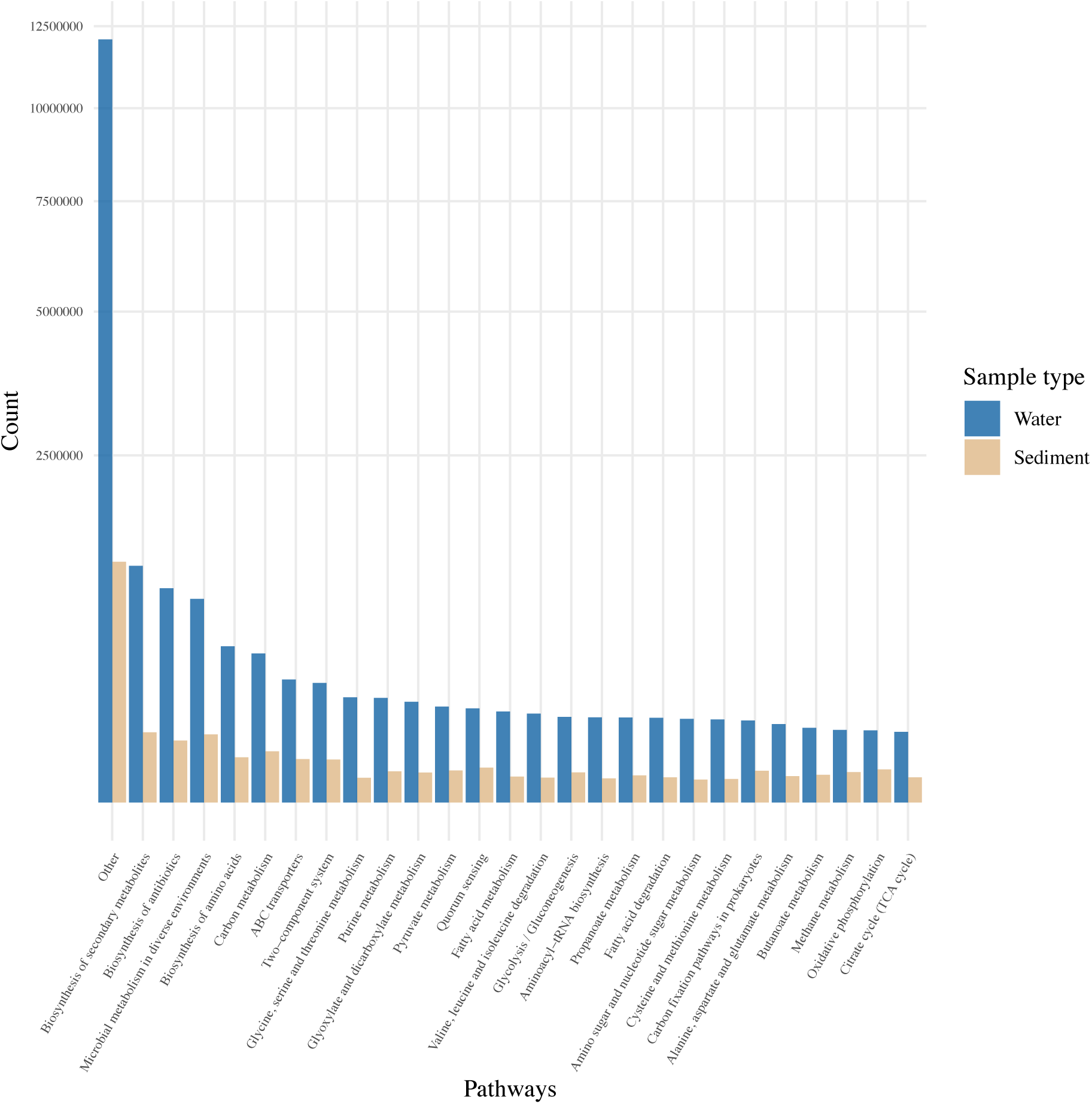
Distribution of proteins associated to KEGG pathways from the plass protein assemblies. The 20 most abundant pathways for both sediment and water are displayed here.

Amongst less abundant but still expressed pathways, we also find carbon, methane and nitrogen metabolism, as well as photosynthesis (Supplementary Fig 3). Additionally, the most abundant pathways associated with antibiotics were related to monobactam, streptomycin and vancomycin. (Supplementary Fig 4.).

**Figure4:**
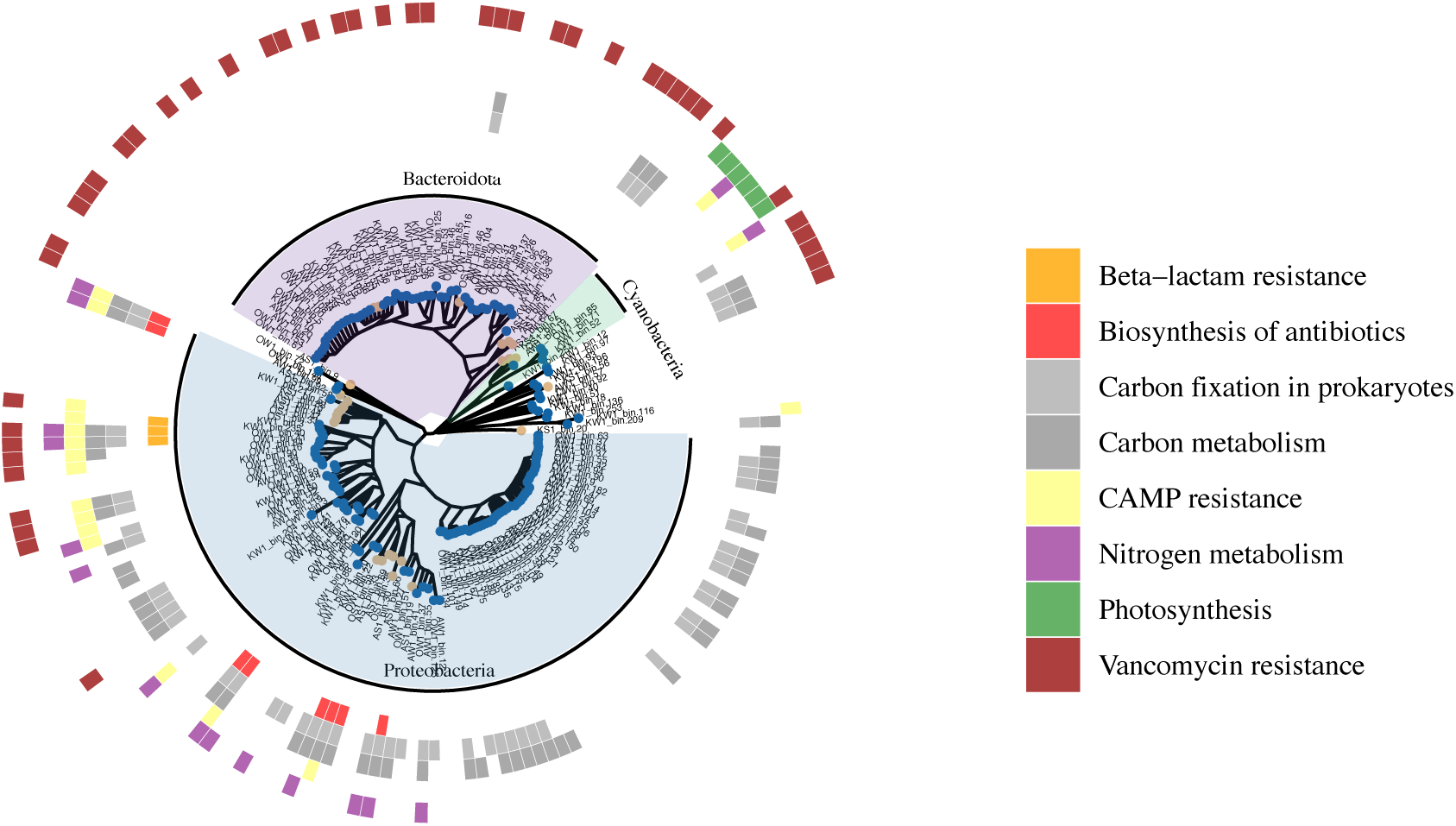
phylogenetic tree for the 178 bacteria recovered from the metagenome assemblies. The tree was generated with pplacer and plotted with ggtree. In blue are the bacteria recovered from the water samples and in brown from the sediment samples. The outer circles represent selected pathways that are present in the assemblies.

#### Metagenome-assembled genomes

A total of 782 genome bins were recovered. Of those, 193 presented more than 50% completeness and less than 15% contamination according to the checkm results. One hundred seventy-eight of those were bacterial genomes, while 15 were classified as archaea.

Twenty-eight of the 178 bacterial genomes had a >95% match to an already published genome. The remaining 150 are either new strains or new species. All genomes were classified to at least the Phylum level. More than half of the recovered Bacteria were Proteobacteria, Bacteroidetes and Cyanobacteria (Fig. 4).

The recovered archaeal genomes all came from water samples. Of the 15, 7 were classified as known archaea from the order Poseidoniales, and eight are newly discovered archaeal species, all putatively placed in the Poseidoniaceae family.

## Discussion

Bacterial communities from Indian Ocean reefs – and coastal waters – are critically understudied. Here we present a catalogue of the microorganisms present in the upper sediment layer and the water near coral reefs, as well as a catalogue of putative proteins and functions of said microorganisms.

This study presents – to the extent of our knowledge – the first metagenomes taken from the coastline of the West Indian Ocean, and offers a baseline for much needed further work on conservation and monitoring of the West Indian ocean coasts. We present and publish a catalogue of 12 million putative proteins and 193 draft genomes, including 175 previously unpublished bacteria. While it had been valuable also to investigate the coral microbiome itself, our study presents a solid baseline for monitoring water quality which we hypothesise may be a good proxy for coral health. Indeed, while physicochemical properties of coastal waters have remained stable in the region regardless of pollution status, bacterial communities may not be.

The taxonomic distribution of bacterial species, as presented in this study, while diverse, is consistent with previously published coastal metabarcoding studies (Kelly et al., 2014). The presence of *Vibrio* in the water samples is worrying, especially given the presence of *Vibrio coralliilyticus* and *Vibrio owensii*. The former may be commensal at some water temperatures, but the pathogenicity of both species is well documented (Gibbin et al., 2019; Ushijima, Smith, Aeby, & Callahan, 2012).

Even though there were no obvious differences in the distribution of phyla between sites and sample types, variations were observed in the abundances of three ecologically important genera. *Candidatus* Pelagibacter, *Prochlorococcus* and *Synechococcus*, which are considered to be ubiquitous in marine environments (Biller, Berube, Lindell, & Chisholm, 2015; Morris et al., 2002; Ruffing, Jensen, & Strickland, 2016), had higher abundances in water at the Kisite-Mpunguti site. *Candidatus* Pelagibacter is believed to be the most successful clade of organisms on Earth and, although a heterotroph, it is known to thrive at the low nutrient concentrations typical of open ocean conditions (Zhao et al., 2017). Overall, members of the *Alphaproteobacteria* class are known to have higher relative abundance in habitats with higher coral cover than in nutrient-rich algae-dominated habitats. Besides *Candidatus* Pelagibacter, no differences were found in the distribution of genera from the class *Alphaproteobacteria*, between sites or samples. On the other hand, members of *Synechococcus* and *Prochlorococcus* are cyanobacteria that are considered the most important primary producers in the tropical oceans, responsible for a large percentage of the photosynthetic production of oxygen (Biller et al., 2015; Kim et al., 2018; Waterbury, Watson, Guillard, & Brand, 1979). Being autotrophs, members of these genera are usually found in great abundance in ocean zones low in nutrients (Dinsdale et al., 2008; Kelly et al., 2014). Also, cyanobacteria have high adaptive capacities for nutrients and light harvesting, a competitive advantage over other marine microorganisms (Louati et al., 2015). As such, *Synechococcus* and *Prochlorococcus* may have outcompeted the other microbes at the Kuruwitu site where its distance from the coast, settlement and increased human activities, may have had limited nutrients essential for their growth.

The main strength of metagenomics over metabarcoding is the insight into function, provided by the protein assembly as well as the binning. We showed that the water samples the presence of many antibiotic-related pathways, for both biosynthesis and resistance. It is particularly striking that the majority of Metagenome-Assembled-Genomes (MAGs) from the *Bacteroidetes* phylum indicated the presence of the vancomycin resistance pathway. While surprising at first given vancomycin is not considered a first-line therapy antibiotic, it may be explained by the use of avoparcin as a food supplement in agriculture. Antibiotic resistant bacteria have been observed widely in aquatic environments following antibiotic contamination from wastewater treatment plants or agricultural runoffs (Schmieder & Edwards, 2012). Avoparcin is a glycopeptide antibiotic that is chemically very similar to vancomycin; there have been earlier concerns about the use of avoparcin in agriculture in various countries as well as Kenya (Bager, Madsen, Christensen, & Aarestrup, 1997; Nilsson, 2012; Raphael, Sam, Anne, Peter, & Samuel, 2017), and it would be reasonable to think it would be the cause of vancomycin resistance gene clusters in coastal waters.

Most assembled proteobacteria showed signs of being autotrophic, presenting carbon fixation and metabolism pathways. Some also seemed to able to fix nitrogen. The cyanobacteria retrieved from the sediment layer exhibited photosynthetic pathways. These organisms may contribute a great deal of nutrient exchange in the whole ecosystem, even though they are not living in direct symbiosis with corals, their contribution cannot be ruled insignificant. Lastly, quorum sensing was found to be one of the most abundant pathways. While relatively little is known about it, bacteria use quorum sensing as a way to chemically communicate with each other, which has its importance in nutrient cycling across and within microbial communities (DeAngelis, Lindow, & Firestone, 2008).

The difference in taxonomic distribution between the read-level classification and the genome binning is quite striking in a few aspects. Almost no draft genomes from the sediment samples passed the threshold for draft genomes of acceptable quality, and this could be explained by a combination of factor and biases at different levels of the experiment, leading to assemblies of poorer quality. All sediment samples resulted in more prominent and more fragmented assemblies (Supplementary table 3) than the water samples. Metagenomes are notoriously tricky to assemble, due to their variable genome coverage and non-clonal nature (Breitwieser, Lu, & Salzberg, 2019), and additional factors such as unusual GC content or repeat-rich genomes may have played a role. The total size of the assemblies may also indicate that the sediment samples are more diverse and complex than their water counterpart, but that diversity and complexity may be poorly represented in our results, due to the incomplete nature of our biological databases and the imperfectability of our algorithms.

In conclusion, the pathways analysis, as well as the annotation of the draft genomes, provide insights into the putative nutrients exchange and other interactions between bacteria and the environment, including corals. Through nitrogen fixation, photosynthesis and carbon metabolism, water and sediment bacteria may prove valuable to nutrient cycling in a healthy reef. Additionally, we hypothesise that coastal bacterial communities are a potential health indicator for reefs, especially regarding antibiotic resistance, potentially from agricultural runoff, as well for opportunistic *Vibrio* pathogens. While metagenomics is expensive and inconvenient to use in a monitoring setting, our dataset may prove valuable in designing more targeted primer-based approaches to detect pollution in coastal communities. It is also possible that – with the advance of long-read portable sequencers such as the oxford nanopore MinION, the cost barrier for field sequencing for monitoring purposes drops dramatically, making metagenomics a viable approach for monitoring coastal communities. Long-read sequencing could also be used for obtaining full-length 16s sequences, which could potentially provide resolution up to the species level.

## Acknowledgements

This work was carried out under research permit from the National Commission for Science, Technology and Innovation (NACOSTI). Kenya Wildlife Service (KWS) provided logistical support and granted permission to work in the marine parks and the National Environmental Management Authority (NEMA) provided access permits.

The authors would like to acknowledge support from Science for Life Laboratory, the National Genomics Infrastructure, NGI, and Uppmax for providing assistance in massive parallel sequencing and computational infrastructure. This work was supported by the Swedish Research Council, grant number 2015-03443_VR.

## Data accessibility

Trimmed sequences were deposited to the European Nucleotide Archive under the study accession PRJEB30838. Scripts and Refining parameters for the genome binning are available at https://osf.io/5fzqu/

## Author contributions

SW designed the study, conducted field sampling and laboratory processing, drafted and revised the manuscript.

HG designed the study, performed the bioinformatics analyses and drafted the manuscript. EV, OKL, NW, EBR, AMD, SV helped design the study and edited the manuscript.

