## Supplementary materials for "Molecular Ecology of Coral Reef Microorganisms in the Western Indian Ocean coast of Kenya"

**Supplementary Table 1**

Measurements of physicochemical characteristics of three coral reefs. Measurements in all sites were completed within 5 days and were done in situ using a portable multiprobe water quality meter. Sites were chosen considering typical prevalent human activities experienced along the Kenya coast of the Western Indian Ocean.

Variable	Kisite	Mombasa	Malindi

Temperature (°C)	28.2	28.6	29.8

pH (unit)	8.0	8.1	8.0

Dissolved Oxygen (ppm)	8.5	8.3	8.2

Salinity (ppt)	30.1	30.1	27.6

Although observed, differences in nutrients concentrations between the study sites were not statistically significant (Supplementary table 2). Consistently the exploited site at Malindi recorded the highest nutrient concentrations while, save for nitrites, Kisite-Mpunguti, the baseline site, recorded the lowest concentrations. Measurements did not vary much across the sites save for temperature and salinity recorded in Malindi that seemed at variance with the other sites: temperature recordings across the study sites ranged between 28.2°C (Kisite-Mpunguti) and 29.8°C (Malindi), while salinity was estimated as 27.6 ppt at Malindi compared to 30.1 ppt in both Mombasa and Kisite-Mpunguti. Dissolved oxygen and pH varied only slightly between the study sites.

**Supplementary table 2**

Nutrient levels measurements of coral reefs in three study sites. Samples were transported to the laboratory on ice and refrigerated before nutrient levels tests were performed within a week. Between-site comparison were tested by analysis of variance (ANOVA).

Nutrient	Kisite	Mombasa	Malindi	*P*

NITRATES (NO_3_^-^ -N)

(mg/L ± SD)	0.002 ± 0.003	0.046 ± 0.029	0.138 ± 0.096	0.068

NITRITES (NO_2_^-^ -N)

(mg/L ± SD)	0.063 ± 0.019	0.059 ± 0.028	0.076 ± 0.005	0.552

AMMONIMIUM (NH_4_^+^ -N)

(mg/L ± SD)	0.035 ± 0.006	0.040 ± 0.006	0.040 ± 0.07	0.572

PHOSPHATES (PO_4_^3-^ -P)

(mg/L ± SD)	0.014 ± 0.013	0.025 ± 0.008	0.041 ± 0.002	0.345

**Supplementary Figure 1**

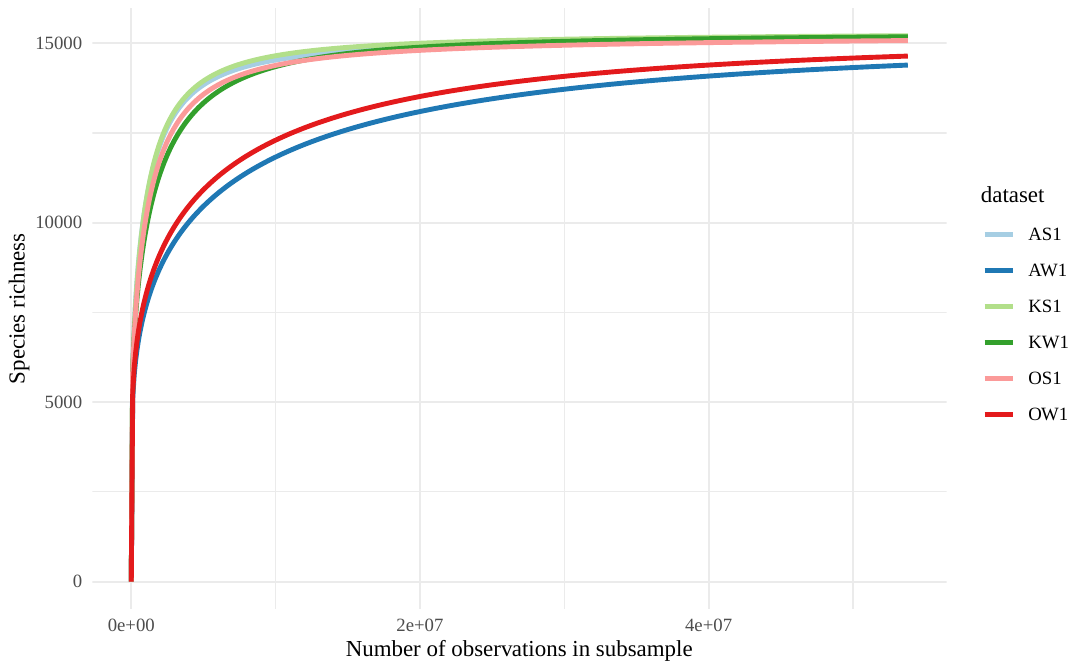

Estimated bacterial richness for all samples. Bacterial richness was estimated from the classified bacteria using the kraken2 results. Datasets are colour coded by location: blue for Malindi, green for Kisite and Red for Mombasa. In the legend S stands for sediment samples, and W for water. The species richness (y axis) represent the distinct number of bacterial species found in the kraken2 results, in function of the total number of observation (x axis).

**Supplementary Figure 2**

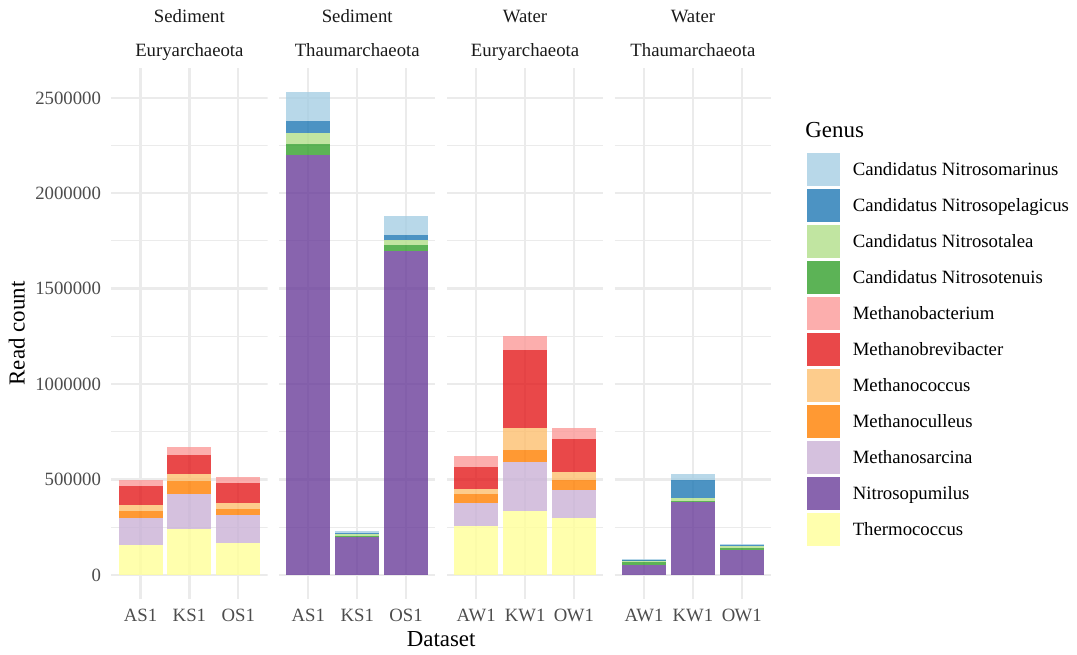

Distribution of classified archaeal reads across datasets, sample types and phyla. On the x axis, A stands for the Malindi sampling site, K for Kisite and O for Mombasa. The Thaumarchaeota, mainly *Nitrosopumilus*, dominate the sediment samples, while Euryarchaeota are more abundant in the water samples.

**Supplementary Figure 3**

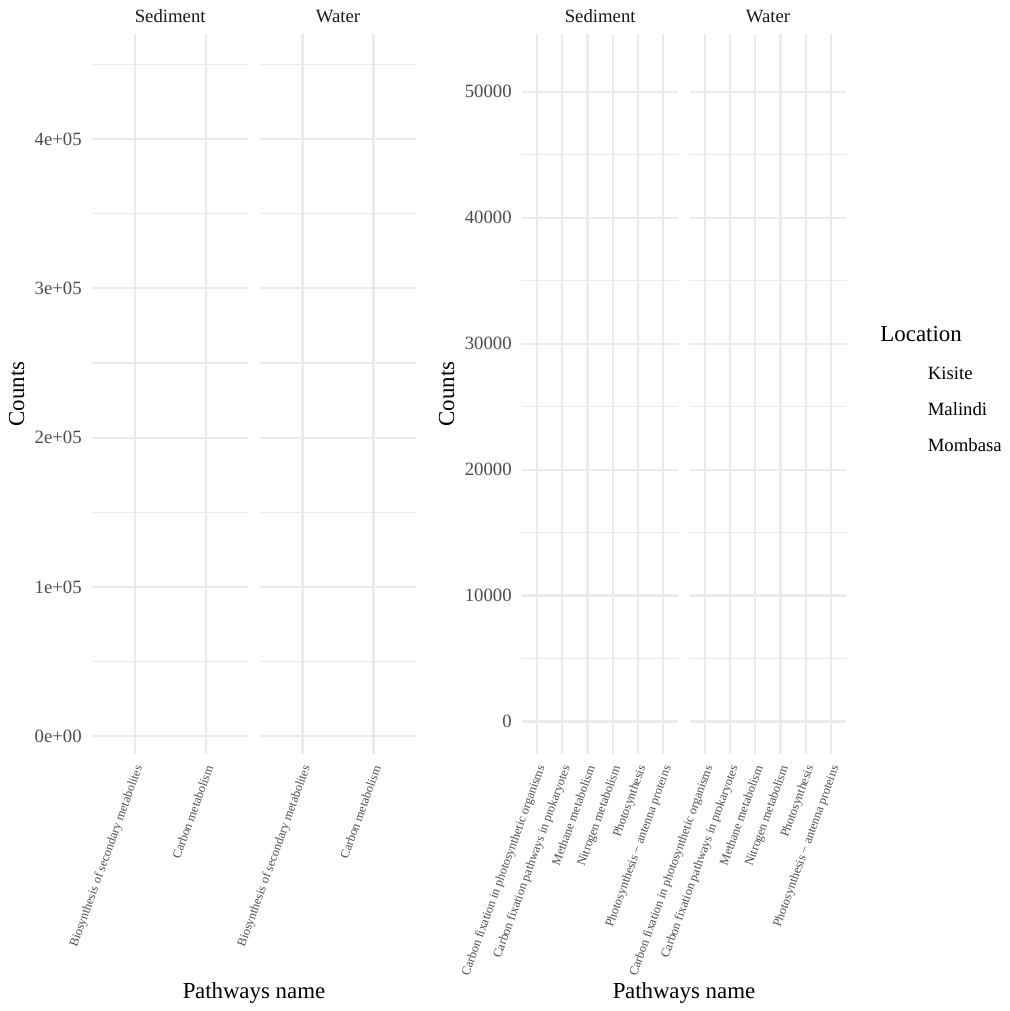

Protein counts for selected pathways, annotated from the protein assemblies. To the left, two of the most abundant pathways, Biosynthesis of secondary metabolites and carbon metabolsim. To the right, less abundant, but still expressed pathways.

**Supplementary Figure 4**

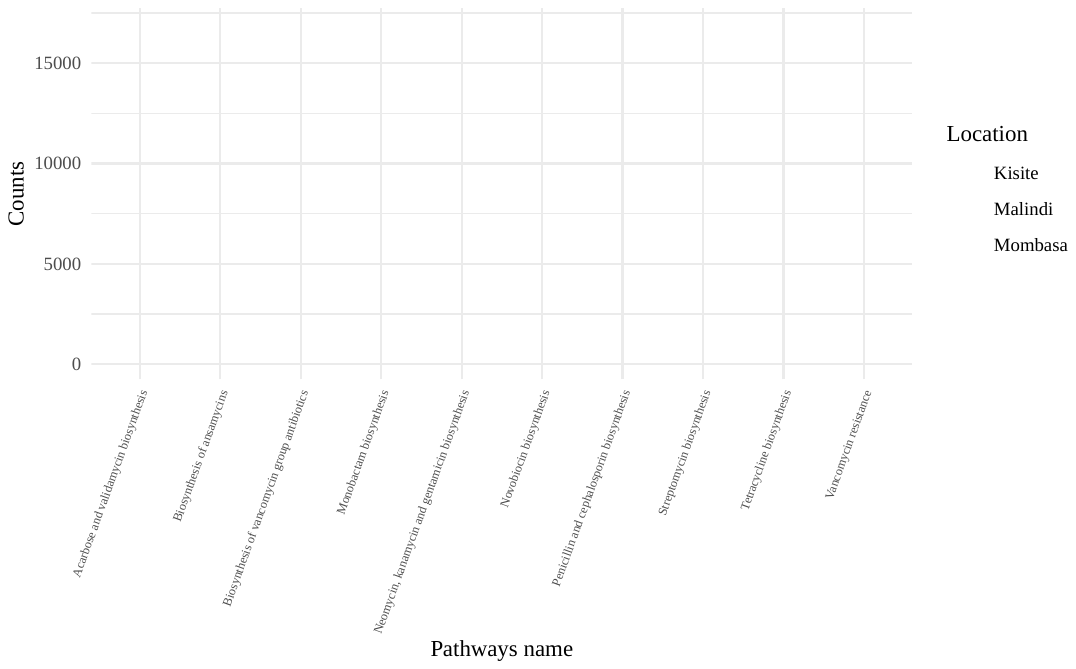

Protein counts for antibiotic-related pathways found in the protein assemblies.

**Supplementary table 3**

| Sample | Location | Sample type | Total length (Mbp) | N. contigs | Longest contig | N50 |
| --- | --- | --- | --- | --- | --- | --- |
| AS1 | Malindi | Sediment | 4255.7 | 4 975 239 | 145 965 | 1117 |
| AW1 | Malindi | Water | 1326.7 | 1 428 285 | 531 348 | 1344 |
| KS1 | Kisite | Sediment | 4454.8 | 5 717 323 | 152 291 | 936 |
| KW1 | Kisite | Water | 5866.7 | 6 784 570 | 178 686 | 1130 |
| OS1 | Mombasa | Sediment | 3895.2 | 4 513 177 | 185 820 | 1140 |
| OW1 | Mombasa | Water | 1493.2 | 1 614 522 | 761 566 | 1396 |

Assembly statistics for all the samples. The sediment samples consistently produced smaller contigs than their water conterparts.
